# Investigation of the mechanism of accelerated biodegradation of *Paracoccus-KDSPL-02*

**DOI:** 10.1101/2024.05.06.592715

**Authors:** Peng Wang, Shanxiang Xu, Chen Shen, Jiewen Ma, Feiyu Cheng, Jingyu Liu

**Author notes:** Correspondence (Peng Wang).

## Abstract

*Paracoccus-KDSPL-02* can accelerate to degrade penicillin G under light remain poorly understood, largely due to the lack of high-throughput genome engineering tools. Firstly, this study sequenced the genome of *Paracoccus-KDSPL-02* and mined the genes that might be involved, and in order to understand in detail whether the expression of the mined genes changed during light. Further, for genes with altering transcriptional levels under light, this study obtained PROKKA_01468 which a photoreceptor protein in *Paracoccus-KDSPL-02*. In the end, for validating the function of PROKKA_01468, this study knocked down the sequence of the PROKKA_01468 by applying gene editing system, and the knockdown strain showed significant change in the rate of degradation of phenylacetic acid, which is the intermediate product of penicillin G degradation, by light compared with darkness, so that the PROKKA_01468 is the most effective photoreceptor protein in *Paracoccus-KDSPL-02*.

**Synopsis:** This research elucidates a molecular mechanism capable of accelerating penicillin G degradation in wastewater, with significant implications for environmental science.

## Introduction

In recent years, with increasing usage, penicillin G contamination has been widely detected in surface water, groundwater, sewage water, and sometimes in drinking water^[1-3]^. Penicillin G and its intermediates could cause potential secondary water pollution^[4,5]^. There are various degradation methods for penicillin G, such as physical degradation, chemical degradation, and biodegradation, among which biodegradation has attracted much attention because of its green, environmental, friendly, and efficient advantages^[6,7]^. Isolation of a strain of *Paracoccus-KDSPL-02* from the sludge of a pharmaceutical factory, which is highly efficient for the degradation of penicillin G^[8]^. The strain has a wide range of degradation capabilities and can biodegrade a variety of compounds^[11]^. It also has the ability to accelerate the degradation of some common and easily degradable compounds ^[12-20]^.

Based on previous studies, it was shown that *Paracoccus-KDSPL-02* accelerates the rate of degradation of penicillin G in the presence of light^[8-10]^. However, the molecular mechanism by which it accelerates the degradation of penicillin G is unclear. Therefore, the whole genome of *Paracoccus-KDSPL-02* needs to be sequenced to mine the relevant genes. Exploration of this mechanism has usually taken the form of speculation about energy changes within the bacterium and changes in protein transcript levels in vivo in response to light^[21-26]^. Further information on photoreceptor proteins can be obtained by analysing them at the transcriptomic level, and changes in NADPH as well as ATP that occur during light exposure can be detected at the same time.

However, this method is not able to visualize the mechanism, It is only possible to show that the expression of the corresponding genes changes in the light, but it is not possible to analysis their specific function. so more effective methods need to be explored. In recent years, gene editing technology has become a powerful tool for investigating the function of genes by knocking out genes and suppressing their expression. Clustered regularly interspaced short palindromic repeats (CRISPR) and CRISPR-associated (Cas) system, which is an RNA guided immune system in many bacteria and archaea, has been extensively studied and harnessed as highly efficient genome editing tools for various microorganism^[27]^. Since 2015, successes have also been reported for genome editing in microorganism using Type II CRISPR-Cas9 system derived from *Streptococcus pyogenes*^[28-30]^. In CRISPR-Cas9 system, the complex of Cas9 and guide RNA (gRNA) can recognize the target sequence containing a specific protospacer-adjacent motif (PAM) “NGG” (N represents any nucleotide) located immediately downstream of the protospacer, and then induces a double-strand breakage.

This study obtained the annotation of *Paracoccus*-related genes by sequencing the genome of *Paracoccus-KDSPL-02* strains, after which This study further mined the sequences related to photoreceptor proteins by sequencing the transcriptome of *Paracoccus-KDSPL-02* after treatment under light. This study further investigated the changes of NADPH and ATP in *Paracoccus-KDSPL-02* on the basis of its gene expression level and metabolic pathway level, and established CRISPR-Cas gene editing system in *Paracoccus-KDSPL-02* for the first time to knock down the photoreceptor protein sequences, and observed the changes of the rate of accelerated degradation of penicillin G and its intermediates in the light-exposed knockout strains. This study also observed the rate change of the knockout strain for accelerated degradation of penicillin G and its intermediates under light exposure, and further derived the molecular mechanism of the accelerated degradation of penicillin G mineralisation in *Paracoccus-KDSPL-02* under light.

## Results

### Mining of photoreceptor proteins by transcriptional analysis in *Paracoccus-KDSPL-02*

In previous research, laboratory found a significant acceleration of penicillin G degradation by *Paracoccus-KDSPL-02* in the exposure of light. It is well known that prokaryotic organisms of the genus Coccidioides are generally not light-sensitive because they do not possess light-sensing organelles such as chloroplasts, so this study hypothesized that there are genetic elements in the genome of *Paracoccus-KDSPL-02* that can sense the light response, and therefore, this study proposed to use transcriptome sequencing in the present study to investigate the changes in the transcript levels of the penicillin G degradation which behaviour of *Paracoccus-KDSPL-02* in the light and the dark conditions, respectively.

The number and distribution of the gene expression value in samples have certain differences. The correlation of gene expression represents the repeatability of biological experiments and can also prove the reliability of subsequent differential analysis. The more similar the sample, the closer the similarity index is to 1 (Fig. 2A). Principal component analysis (PCA) reflects the differences of multiple sets of data on the coordinate axes. The more similar the sample composition is, the closer the distance is reflected in the PCA graph (Fig. 2B). PCA showed that the distribution of the experimental treatments was substantially different. It can be seen from the figure that there are differences among the 6 samples, and the differences are significant. And the significant difference in gene expression between light and dark conditions was quite significant. COG analysis showed that the functional groups with the most differential genes were Amino acid transport and metabolism; Inorganic ion transport and metabolism; Energy production and conversion. The main gene types are nitrate reductase alpha/beta/gamma/delta subunit; ABC-type amino acid transport; nitrate/nitrite transporter; Aminopeptidase; Acetone carboxylase, gamma/beta subunit; Transcriptional regulator; Outer membrane receptor proteins, mostly Fe transport; Cytochrome C and so on. The significant difference in one of the outer membrane receptor proteins suggests that one or more of the unvalidated photoreceptor proteins obtained through genome mining exist to sense the light response, which in turn creates functional motif differences.

**Fig. 1.**
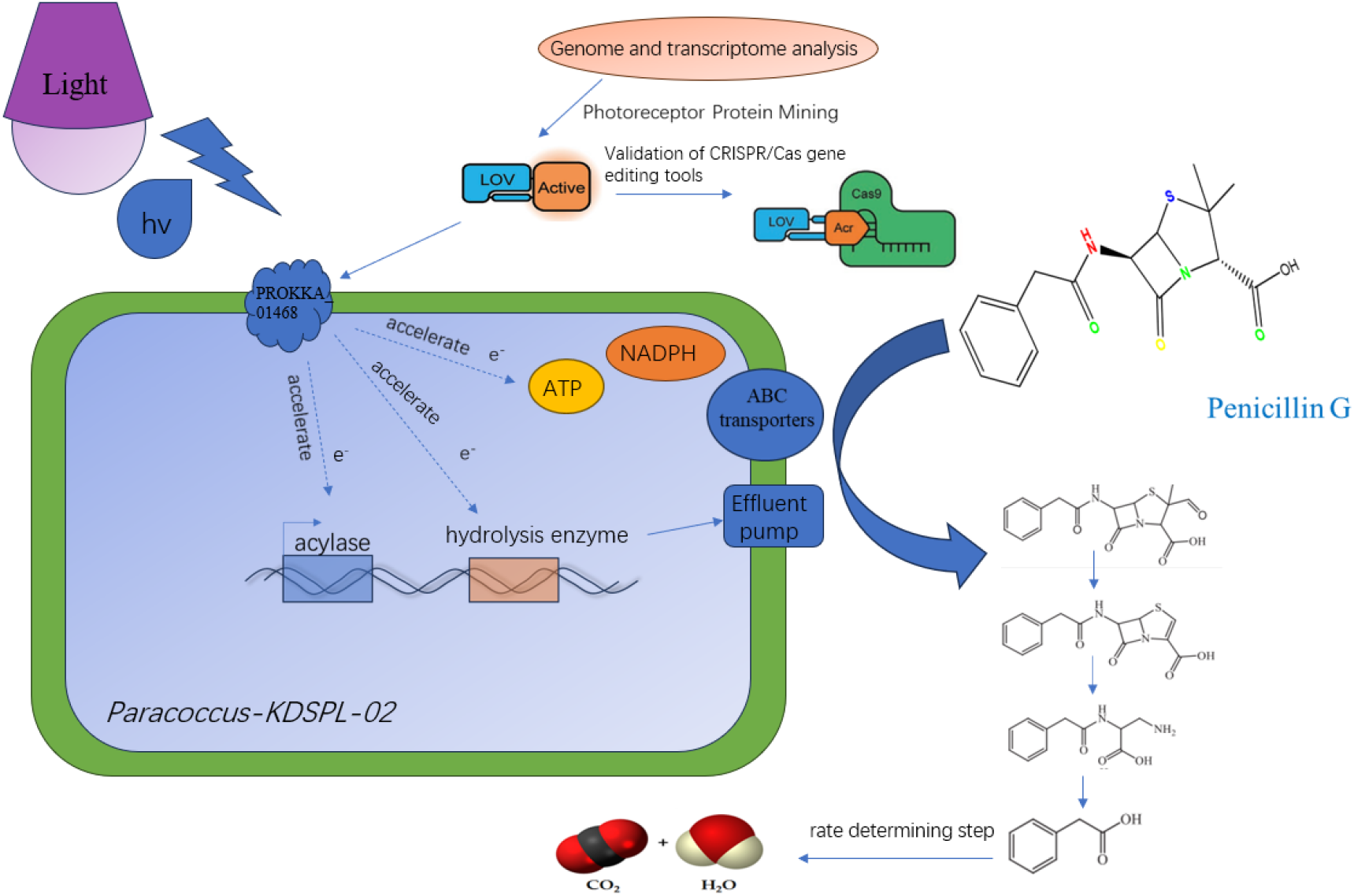
Possible mechanism of accelerated penicillin G degradation under light exposure in *Paracoccus-KDSPL-02*.

**Fig. 2.**
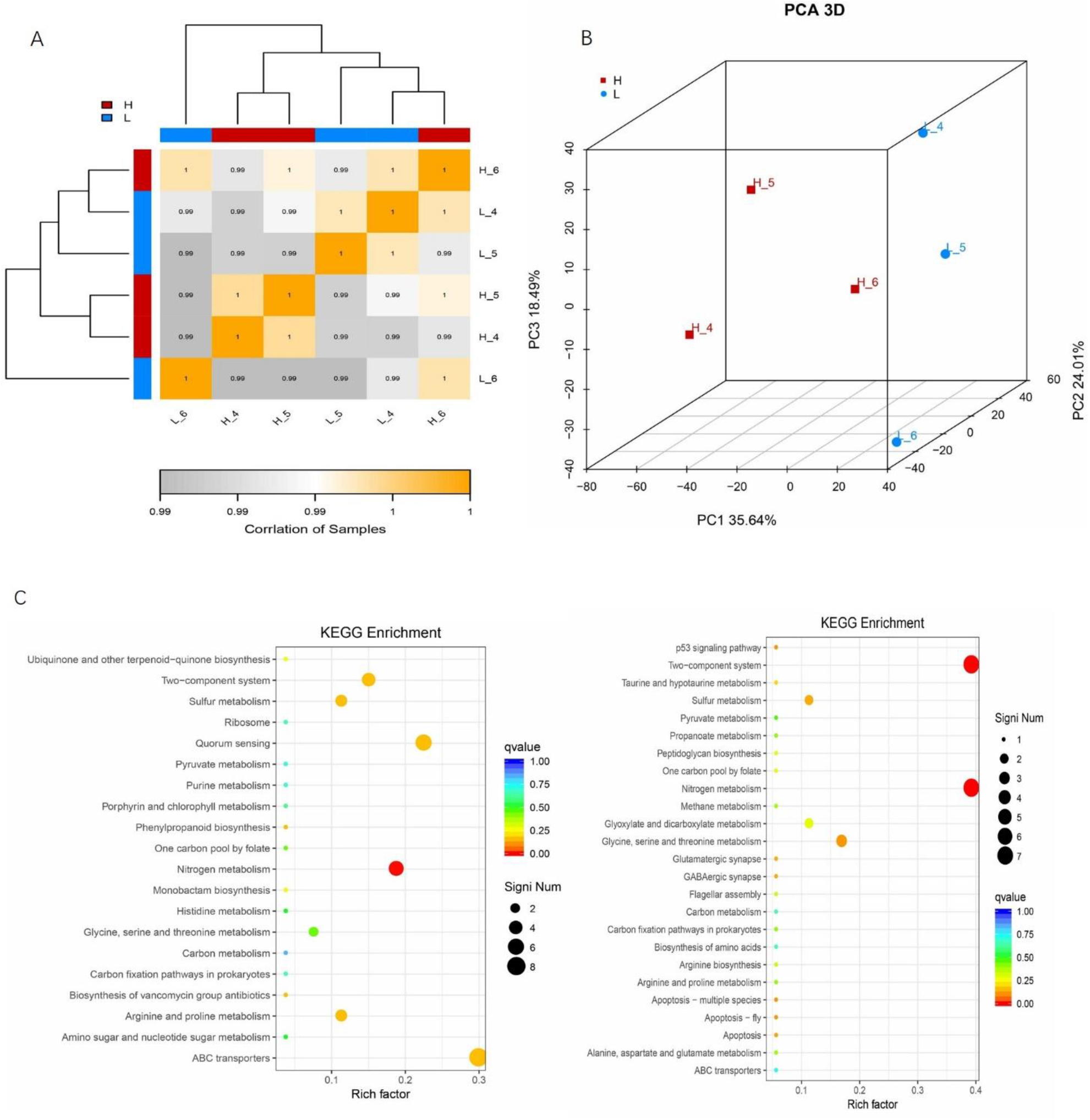
Transcriptional analysis of *Paracoccus*.*KDSPL-02* (A) Sample correlation analysis Heatmap;(B) Principal component analysis (PCA) diagram (C) KEGG Significantly enriched functional scatter plot.

KEGG analysis showed that the DEGs were remarkably enriched in Two–component system (ko02020); Glycine, serine and threonine metabolism (ko00260); ABC transporters (ko02010); Carbon metabolism (ko01200); Nitrogen metabolism (ko00910); Sulfur metabolism (ko00920); Quorum sensing (ko02024); Arginine and proline metabolism (ko00330). One carbon pool by folate (ko00670); Carbon fixation pathways in prokaryotes (ko00720); Pyruvate metabolism (ko00620) and Biosynthesis of amino acids (ko01230) are also involved (Fig. 2C).

Differentially expressed genes in light response can be identified using transcriptome data to gain insight into molecular mechanisms. The analysis of differentially expressed genes is based on the comparison between each two groups of samples, and the final result is that the treatment group is up-regulated or down-regulated relative to the control group. This study found significant changes in the results of the three parallel comparisons, In L 4 vs H 4, a total of 98 genes were significantly changed, of which 43 genes were significantly up-regulated and 55 genes were significantly down-regulated; In L 5 vs H 5, a total of 86 genes were significantly changed, of which 52 genes were significantly up-regulated and 34 genes This were significantly down-regulated. In L 6 vs H 6, a total of 32 genes were significantly changed, of which 15 genes were significantly up-regulated and 17 genes were significantly down-regulated. The L vs H group was the data obtained by DESeq2 analysis of repeated experiments (Fig. 3).

**Fig. 3.**
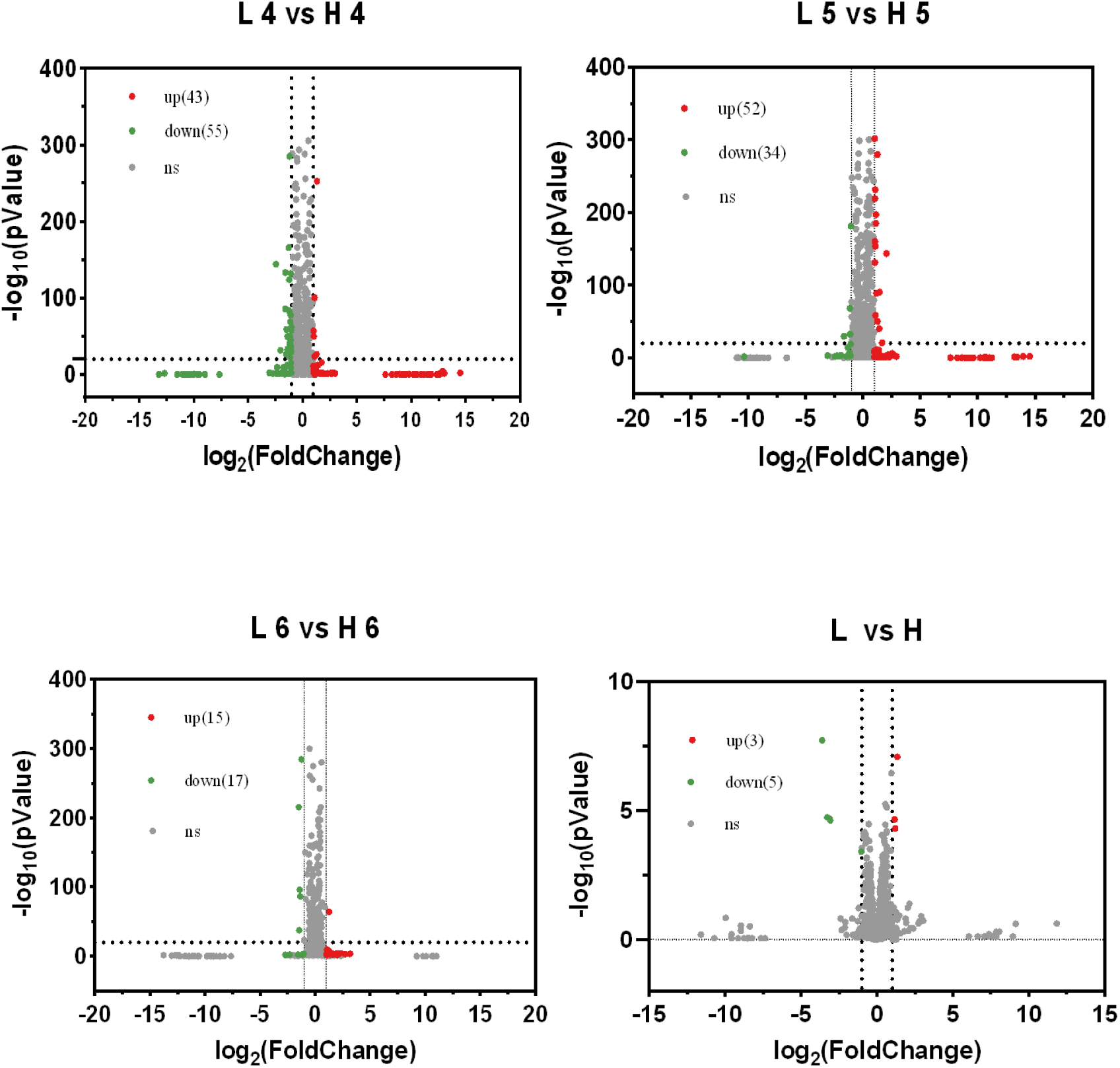
Comparison group expression difference volcano diagram

Metabolic pathway analyses showed significant changes in ABC transporters in this process and a significant population effect, so that photoreceptor proteins in *Paracoccus-KDSPL-02* could be transcribed and produce a range of responses in response to light exposure. The significant change of ABC transporter indicates that the energy changes of ATP and NADPH occurred in *Paracoccus-KDSPL-02*, so the energy changes occurred in the body can be explored later.

### Analytical mining of structural domains of photoreceptor proteins

As shown in the transcriptional analysis, the transcriptional levels of several genes in *paracoccus* sp. KDSPL-02 can be modulated by light. The modulated gene mainly distributed in ABC-transport and metabolic related proteins. These results revealed that there must be one or more functional protein in *Paracoccus* sp. KDSPL-02 that can response light. There has been evidence that the light response of *Bacillus* is due to the light-sensitive proteins ^[29,30]^. A homologous protein (PROKKA_01468 from *Paracoccus-KDSPL-02*) of blue-light photoreceptor in *Bacillus* (GenBank: AAC00382) was found through screening the genome of *Paracoccus-KDSPL-02* as shown in Table1. The domain of PROKKA_01468 contained PAS and PAC that is extremely similar with the blue light receptor proteins in *Bacillus* through SMART domain analysis. We proposed that PROKKA_01468 was responsed for perceiving light signals, triggering light-response pathways and activating transcription. For the functional validation of this photoreceptor protein, the CRISPR/Cas gene editing system was established and used for the knockout of this photoreceptor protein gene in *Paracoccus* sp. KDSPL-02 to investigate whether PROKKA_01468 can response light and the effect of light exposure on the degradation rate of penicillin G and its intermediates.

**Table 1.** Light sensitive protein in KDSPL-02.

| Gene ID | Query Length | Subject |
| --- | --- | --- |
| PROKKA_01468 | 327 | Blue-light-activated histidine kinase 1 |
| PROKKA_01710 | 679 | Blue-light-activated protein |
| PROKKA_02852 | 626 | Blue-light-activated protein |
| PROKKA_03479 | 188 | Methylamine dehydrogenase light chain precursor |
| PROKKA_04256 | 219 | Light-independent protochlorophyllide reductase iron-sulfur ATP-binding protein |
| PROKKA_05001 | 364 | Blue-light-activated histidine kinase |

### Harnessing the endogenous Type I-C CRISPR-Cas system for genome editing in *Paracoccus-KDSPL-02*

As previously indicated, the endogenous type I-C CRISPR-Cas system of *Paracoccus-KDSPL-02* exhibits good interference activity against vectors harboring both the correct PAM sequence and the original spacer sequence. The technique exhibits a high level of interference activity against chromosomal-carrying vectors (self-targeting), which results in effective selection on wild-type cells. Next, our study tried to alter the *Paracoccus-KDSPL-02* genome seamlessly using the type I-C CRISPR-Cas method (Fig. 4A).

**Fig. 4.**
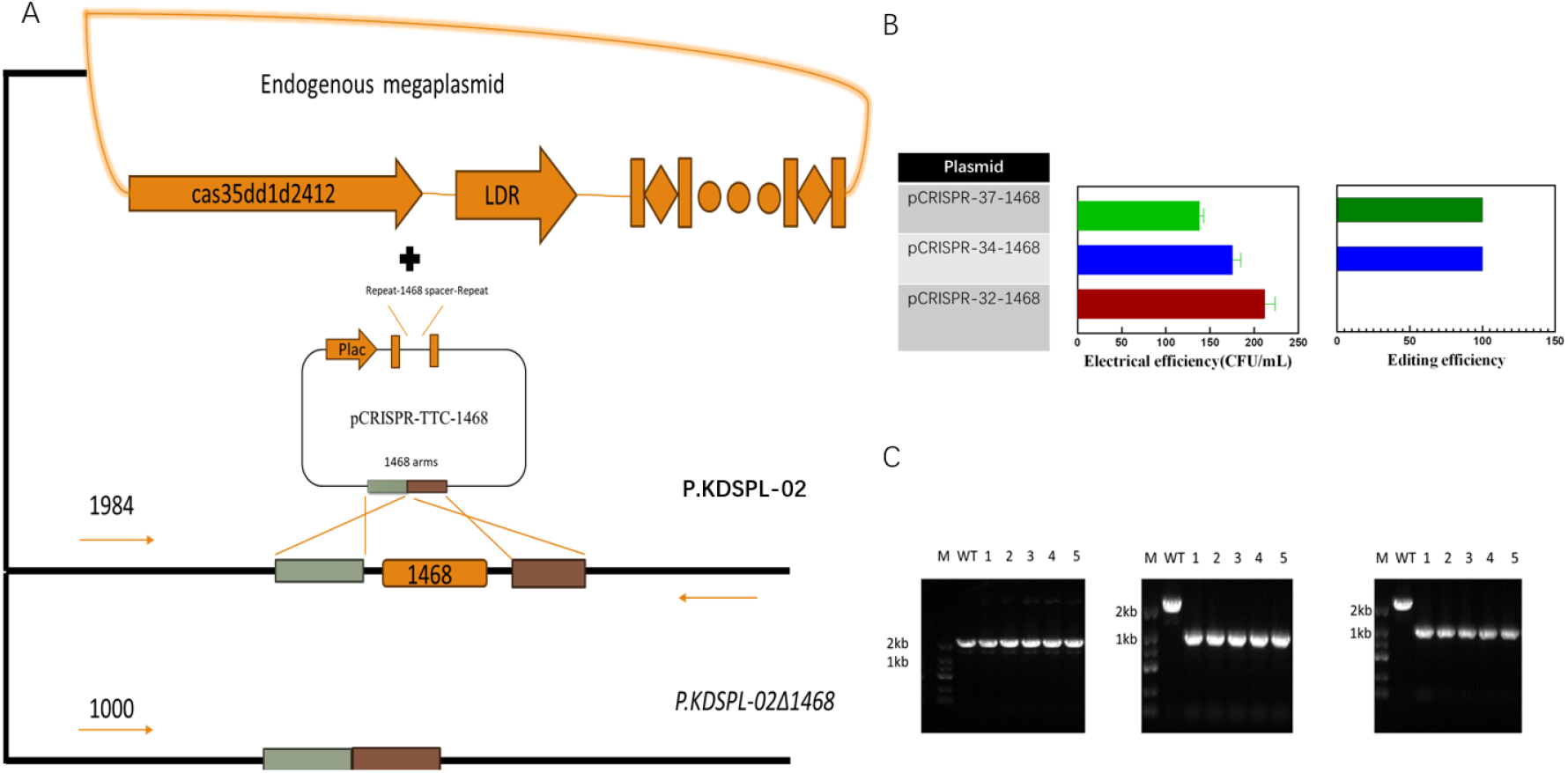
Harnessing the endogenous Type I-C CRISPR-Cas system for genome editing in *Paracoccus-KDSPL-02* (A) Schematic representation of the knockdown of the endogenous CRISPR/Cas system in *Paracoccus-KDSPL-02* (B) Effect of gRNA length on transformation efficiency and editing efficiency. (C) Agarose gel electrophoresis of different lengths of gRNA knockdown efficiency

The two components of the genome editing plasmid are a pair of homology arms for gene editing and a synthetic CRISPR sounding cassette for self-targeting. The process of creating the synthetic CRISPR expression cassette involved imitating the native gene. In order to boost editing efficiency and decrease the self-targeting activity of the endogenous type I-C CRISPR-Cas system, a lactose-inducible promoter was employed in this study to build the endogenous CRISPR arrays. A synthetic CRISPR array was transcribed under the direction of a lactose-inducible promoter. Two 32 nt straight repeat sequences and a 34 nt spacer sequence make up the array. Once more, the target gene was chosen as 1468. There are a total of 13 possible PAM sequences for the 984 bp 1468 gene. For the self-targeting location, this study chose a PAM sequence along with its downstream 34 nt original spacer region.

Plasmid pCRISPR-TTC-1468, which has two homology arms (around 500 bp each) and an inducible CRISPR expression cassette After construction, the pCRISPR-TTC-1468 plasmid was transfected. A total transformation efficiency of 150–200 CFU/ml was observed in *Paracoccus-KDSPL-02* (Fig. 4B). In order to induce transcription of synthetic CRISPR arrays, the transformants were obtained and then put to LB liquid media and cultivated onto solid plates containing IPTG. Five colonies were chosen at random, and the 1468 gene was checked for. According to the colony PCR results, every tested colony generated transcripts identical to the Δ1468 genotype, indicating a 100% editing efficiency.

The length of the spacer gRNA was optimized in this chapter in order to investigate the editing efficiency of the *Paracoccus-KDSPL-02* endogenous CRISPR system in more detail and to confirm the impact of spacer length on editing efficiency and transformation efficiency. Because the endogenous CRISPR system’s transformation efficiency is still very low, spacer gRNAs of 32 and 37 nt were set in addition to the 34 nt previously mentioned, and the genome editing and transformation efficiency of the strain were examined using the spacer gRNAs of various lengths. Knockout plasmids with spacers of 32, 34 and 37 nt were created and given the names 32-gRNA-1468, 34-gRNA-1468, and 37-gRNA-1468 in order to investigate the impact of spacer region length. For knockout validation, three plasmids were inserted into the *Paracoccus-KDSPL-02* strain, accordingly. Following induced transcription, five colonies were selected at a time, and ten colonies in total underwent colony PCR for confirmation (Fig 4C). The findings demonstrated that while the knockdown efficiency approached 100%, the transformation efficiency dropped by over 20% when the spacer region gRNA sequence length was 37 nt as opposed to 34 nt. The endogenous CRISPR-Cas system’s editing effectiveness fell as a result of the short spacer, even though the transformation rate remained high, at 32 nt when the knockdown efficiency was 0, appearing in the same band as the wild type. For the endogenous CRISPR-Cas system of *Paracoccus-KDSPL-02*, taking into account both transformation efficiency and editing efficiency, the spacer length of 34 nt is significant for both transformation efficiency and editing efficiency, meaning that 34 nt is the ideal spacer gRNA length for the *Paracoccus-KDSPL-02* strain.

### Characterization of mutant phenotypes

The initial carbonylation reaction on the bis-methyl side chain, followed by cyclization to form a thiazole ring, which yields the by-product Penicillium thiazole acid, and further degradation of Penicillium thiazole acid into phenylacetic acid, which is then mineralized and degraded into CO_2_ and H_2_O, which is also the rate-limiting step of the total degradation process, are the steps that confirm the process of penicillin G degradation by *Paracoccus-KDSPL-02*, as preliminary laboratory work verified the process.

Penicillin G concentration steadily decreased during the degradation process, and as a result, the yield of its intermediates increased. However, the conversion rates of these intermediates— potassium penicillothiazoleate and penicillothiazole acid—were relatively low, and light did not significantly speed up the conversion. However, light did significantly speed up the degradation of penicillin G next degradation product, phenylacetic acid (Fig. 5B). It is important for light to speed up the mineralization of phenylacetic acid into CO_2_ and H_2_O since this process is both a crucial step in the degradation process and a rate-limiting reaction.

**Fig. 5.**
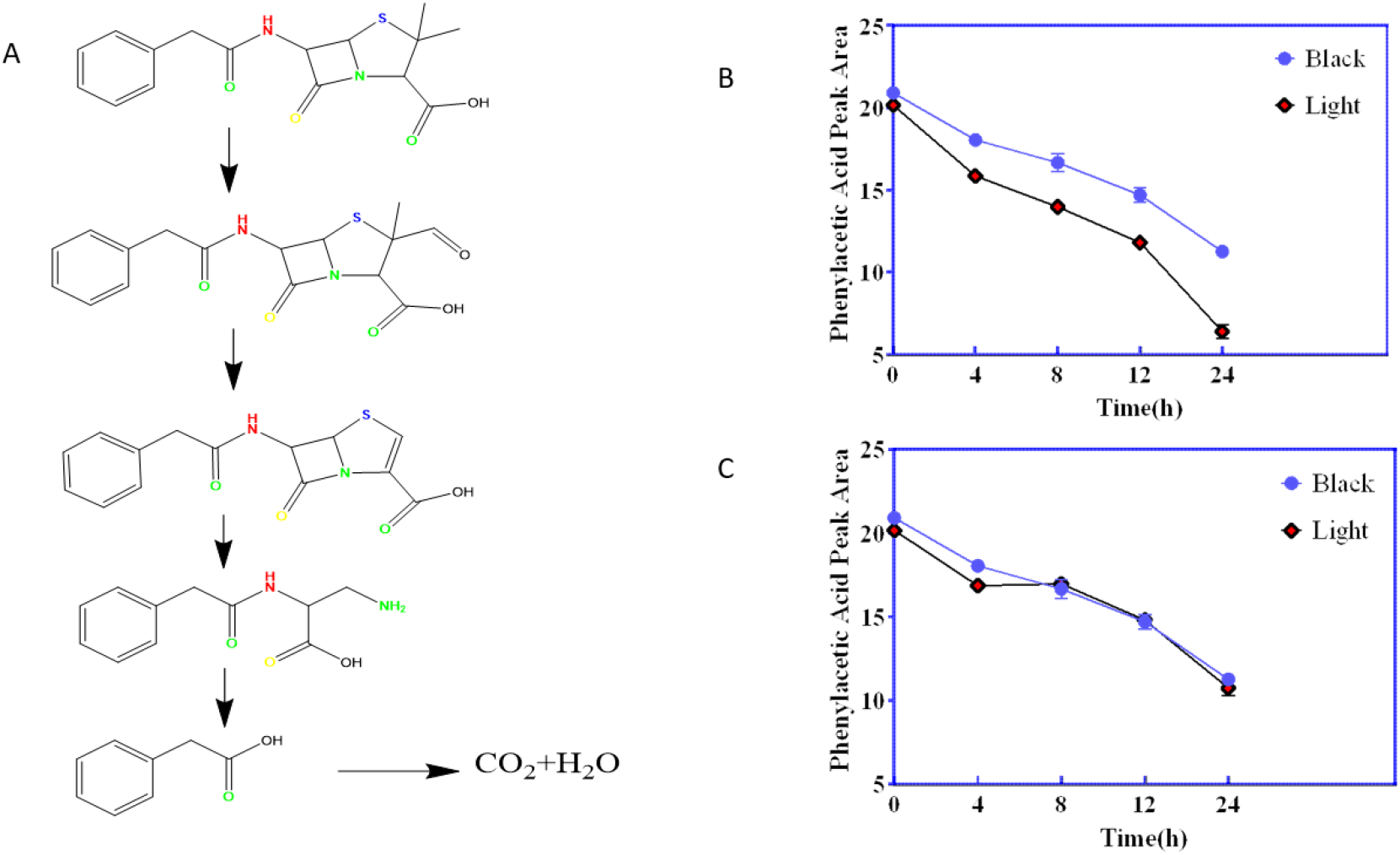
Characterization of mutant phenotypes (A)Degradation of penicillin G in the presence of *Paracoccus-KDSPL-02*. (B)Changes in phenylacetic acid under light and dark in the knockout strain. (C) Changes in phenylacetic acid in wild strains under light and darkness

This study knocked out this gene in *Paracoccus-KDSPL-02* using the experimental method previously described in order to investigate whether the light-accelerating mechanism plays a role in the photosensitive proteins expressed by this gene. Additionally, the degradation process of phenylacetic acid was characterized by HPLC (Fig 5C). The findings demonstrated that, in light circumstances, the knockout strain was less able to mineralize and break down phenylacetic acid, which is an intermediate product of penicillin G. The kinase structure found in the protein structural domain of PROKKA_01468 is thought to be responsible for the bacteria’s ability to detect changes in their surroundings. One of the roles of the kinase structure is to aid in the sensing of external light sources. The mechanism that speeds up the mineralized degradation of phenylacetic acid in the presence of light was lost when the gene was knocked out of *Paracoccus-KDSPL-02*. Additionally, the bacteria were unable to transmit the light-sensing response to a number of light-sensing proteins due to their lack of light-sensing ability.

### General features of *Paracoccus-KDSPL-02* genome

Here, the genome of *Paracoccus-KDSPL-02* was sequenced using the Illumina Hiseq 2500 platform. This study got some genetic prediction information from genomic data. Component analysis mainly includes coding gene prediction, non-coding RNA prediction, repeat sequence analysis, CRISPR sequence prediction and other analysis. The genome of *Paracoccus-KDSPL-02* is linear and does not contain plasmids. Briefly, the sequencing generated 182 reads with mean read length 33522 bp, totaling 5531287 bp. Where the total number of coding genes 4854161 bp. The minimum length is 49bp, the maximum length is 8289 bp. The GC content of the strain is 67.51%. A total of 5397 protein-coding genes, 3 rRNA, and 52 tRNA genes are found in the strain. This complete genomic information provides a clear genetic background for subsequent study of the mechanism of photocatalytic biodegradation in *Paracoccus-KDSPL-02*.

Functional annotation: Among the 5397 predicted genes, 5212(98.06%) genes could be annotated by BLASTN using NCBI Nr databases based on sequence homology. In addition, 2447 (46.04%), 4111 (77.35%), 3707 (69.75%),3873 (72.87%), 4175(78.55%) and 5186 (97.57%) genes could be annotated according to KEGG, COG, GO, SwissProt, PFAM, and TrEMBL databases, respectively. It should be noted that among these genes assigned to Nr database, the top 2 species of matched genes number are *Paracoccus-KDSPL-02* versutus (4054,75.11%), and *Paracoccus-KDSPL-02* (199, 3.69%). Furthermore, 3707 genes could be classified into three Gene Ontology (GO)categories: cellular component, biological process, and molecular function. As shown in the Fig. S1A. KEGG annotation diagram is shown in Fig. S1C. Genes in KEGG annotation mainly focus on carbohydrate metabolism and amino acid metabolism. COG annotation diagram is shown in Fig. S1B.

Through the analysis of the genome sequencing results, it is easy to see that the genome of *Paracoccus-KDSPL-02* has the highest abundance of genes related to metabolic processes, and the process involves more activities related to nucleotides and transcription factors, which suggests that although *Paracoccus-KDSPL-02* does not have endogenous plasmids, many genes in its genome have considerable transcriptional and expression activities. Therefore, the photoreceptor protein can be used to accelerate the degradation of penicillin G. The photoreceptor protein can be applied to other metabolic processes to make the photoreceptor protein function universal.

## Discussion

In this study, the genes that can accelerate the degradation of penicillin G under light in *Paracoccus-KDSPL-02* were mined, and found some genes that can be up-regulated under light by transcriptomic analysis. Combined with the gene annotation in genomics and the analysis of structural domains, this study hypothesized that PROKKA_01468 might be a photosensitive protein that plays the role of light sensing in *Paracoccus-KDSPL-02*. In this experiment, the endogenous CRISPR-Cas system of *Paracoccus-KDSPL-02* was mined and established, and the knockout of the photosensitive protein PROKKA_01468 was achieved, the degradation of penicillin G by the knockout strain was observed to be not significantly accelerated under both light and dark conditions. The key gene for accelerated degradation of penicillin G in *Paracoccus-KDSPL-02* under light conditions is PROKKA_01468, in which there is a LOV light-sensing domain in the sequence of the photoreceptor protein PROKKA_01468, as well as a kinase domain, and the electron channel of the kinase domain is closed under dark conditions, which results in the LOV domain not being able to contribute to the metabolism of a wide range of metabolic processes in *Paracoccus-KDSPL-02*. The LOV domain does not promote a wide range of metabolic processes in *Paracoccus-KDSPL-02*, and inhibits many metabolic processes to a certain extent. Light can open the electron channel of the kinase domain and enable the LOV domain to realize its electron transfer function and participate in metabolic processes, accelerating the metabolic pathway.

Light-sensitive proteins are not commonly found in prokaryotes, and they are mainly found in fungi. Light controls important physiological as well as morphological responses in fungi^[33-35]^. Fungi can use up to 11 photoreceptors and signaling cascades to sense near-ultraviolet, blue and green light. Fungi exert transcriptional regulation at different sites under different light conditions, with blue light receptors exerting transcriptional regulation mainly in the nucleus and green light receptors exerting transcriptional regulation mainly at the cell membrane^[36-38]^. In this study, *Paracoccus-KDSPL-02* can significantly promote the degradation of penicillin G under green light, so This study hypothesized that its photoreceptor protein PROKKA_01468 is a green photoreceptor-related gene on *Paracoccus-KDSPL-02*.

In this study, the genome of *Paracoccus-KDSPL-02* strain was deeply excavated. From the GO classification data of the genomic data, it can be seen that the main classifications of gene functions in *Paracoccus-KDSPL-02* strain can be divided into the following three categories: Biological process, Cellular component, and Molecular function. As can be seen from the abundance values, in the *Paracoccus-KDSPL-02* strain genome, the genes associated with the biological reaction process Therefore, the genes related to light-accelerated penicillin G degradation in *Paracoccus-KDSPL-02* strain are likely to appear in this classification. Most of these genes are related to the relevant metabolic reactions, and the number of genes is large, which needs to be further explored in combination with transcriptomics analysis.

Through genomic analysis, this study posits the existence of gene clusters within the bioprocess compartment, which encode for photoreceptor proteins. These proteins are implicated in the enhancement of penicillin G degradation when exposed to light. For further excavation, this study cultured *Paracoccus-KDSPL-02* strain under light and darkness and sequenced the transcriptome. In the exploration of the genome for ABC transporters, this study identified seventy-three potential target genes. Transcriptome analysis showed that light also promoted the change of ABC transporter. ATP-binding cassette (ABC) proteins are found in all organisms, from bacteria to humans. The ABC transporter family is one of the largest eukaryotic protein families. It’s also found in prokaryotes. Its members play roles in numerous metabolic processes in bacteria by releasing energy for substrate transport across membranes through hydrolysis of ATP^[39]^. The ABC proteins are membrane transporters. ABC regulate cellular levels of endo- and xenobiotics by transporting molecules across cell membranes and are involved in diverse biological processes ^[40]^. As these proteins are linked to resistance, they are often referred to as multidrug resistance proteins. ABC transporters may be involved in the efflux of drug metabolites ^[41]^. The HKs that contribute to QS are part of the two-component signal transduction systems (TCSs). TCS is the main mechanism in bacteria for responding to environmental stimuli and prevails across the entire bacterial kingdom^[42]^. Bacteria use two-component system (TCS) to sense and respond to changes in their environment. Each two-component system consists of a sensor protein-histidine kinase (HK) and a response regulator (RR). The two-component pathway typically enables cells to perceive and respond to stimuli by inducing changes in transcription. Transcriptome analysis indicated that TCS is the key for light sensing in *Paracoccus-KDSPL-02*, which are closely connected to glutamine synthetase, Nitrate reductase subunit, cytochrome C, resulting in an increase related transcription factor^[43]^.At the heart of the TCS signaling pathway are multi-domain sensor histidine kinases that undergo autophosphorylation in response to specific stimuli, triggering downstream signaling cascades^[44]^. Bacterial uptake of amino acids from extracellular space provides cells with compounds that can be used as carbon, nitrogen or energy, which may result in changes in amino acid transport and metabolism. Some studies have shown that some amino acids are decomposed, and the carbon skeleton can be converted into metabolites such as pyruvate through catabolism under stress conditions, affecting amino acid metabolism. The combination of amino acid catabolism and gluconeogenic pathways may be a mechanism for adaptation to light intensity.^[45]^

The ability to sense and respond to environmental cues is essential for adaptation and survival in living organisms. In bacterial systems, exposure to light triggers a conformational shift of proteins into a photoactivated signaling state. This transformation is facilitated by the multidomain sensor histidine kinases integral to the bacterial two-component systems. These kinases initiate autophosphorylation upon specific environmental stimuli, subsequently propagating downstream signaling cascades. The resultant effects are mediated through alterations in enzyme activity, protein-protein interactions, or the modulation of gene expression by functioning as transcription factors. ^[44,46,47]^. Through the established endogenous CRISPR-Cas system based on *Paracoccus-KDSPL-02* endogenous knockout of PROKKA_01468 and subsequent validation by degradation experiments, this study confirmed the conjecture.

Next, for validating the function of the mined photoreceptor proteins, this study set up an endogenous CRISPR-Cas gene editing system in *Paracoccus-KDSPL-02*. Repurposing of endogenous CRISPR-Cas machinery for genetic engineering in *Paracoccus-KDSPL-02* has considerable advantages over heterologous Type II CRISPR-Cas9 system established in this bacterium. First of all, the endogenous CRISPR-Cas system does not need to express the large 4.1 kb Cas9 gene, which could reduce the size of editing vector and decrease the toxicity of this system to the host cells. Thus, the endogenous CRISPR-Cas machinery requires lower demand of plasmid conjugation efficiency. Second, the endogenous CRISPR-Cas machinery employs more compact RNA guide which would simplify the plasmid construction process, especially when multiplexed editing strategies are used. Last, the spacer sequence used for gene targeting in endogenous CRISPR-Cas machinery is much longer than that used in CRISPR-Cas9 system (32– 37 nt vs.20 nt) which could mitigate the potential off-target effect. After completing the knockdown of the PROKKA_01468, this study found that the accelerating effect of the rate-limiting step of penicillin G degradation, which can not accelerate the mineralized degradation of phenylacetic acid under the original light, and PROKKA_01468 is a key protein for sensing the light response in *Paracoccus-KDSPL-02*, which is of very broad application value.

In conclusion, this study obtained functional classification information in *Paracoccus-KDSPL-02* by genomics analysis. In order to investigate genes that changes in expression levels in light, this study applied transcriptional technology and discovered PROKKA_01468 which a highly efficient photosensitive protein. This speculation was verified by the knockout function of the endogenous CRISPR-Cas system in *Paracoccus-KDSPL-02* and the comparison of photodegradation properties of the knockout strain with wild strain. This study investigates the molecular mechanism which PROKKA_01468 accelerates penicillin G degradation in prokaryotic organisms, and further elucidates the importance of photoreceptors for microbial metabolic processes, which can help to guide the development of photoreceptors to control optogenetic tools for gene expression in cells and organisms. It has important value for the development of photo-enzymatic systems.

## Materials and methods

### Bacterial strains and cultivation

Table 1 lists all the strains and plasmids used in this study. *E. coli* DH5α strain (Sangon Biotech) was used for DNA cloning. Transformation of exogenous plasmid systems into *Paracoccus-KDSPL-02* by a gene importer. *E. coli* and *Paracoccus-KDSPL-02* strains were cultured in Luria-Bertani (LB) medium supplemented with 50μg/ml kanamycin (Kan) and or 25μg/ml chloramphenicol (Cm) when required. *E. coli* strain was routinely propagated at 37°C in Luria-Bertani (LB) medium. *Paracoccus-KDSPL-02* strain were routinely propagated at 28~30°C in Luria-Bertani (LB) medium. 0.5mM IsoPropyl b-D-ThioGalactoside(IPTG)for inducing function in the mini-CRISPR System.

### Plasmid construction

To construct CRISPR-Cas9-assisiting plasmids used in *Paracoccus-KDSPL-02*, the plasmid pCas9 and PUC57-sgRNA, which was previously developed in prokaryote, was chosen as the mother vector. For the attempt to delete PROKKA_01468 gene, the sgRNA fused with 20-nt guiding sequence into pCas9 and two homology arms (~500 bp each) were inserted into the EcoRI and BamHI sites of PUC57-sgRNA. In order to use the endogenous CRISPR system of *Paracoccus-KDSPL-02* to knock out, this study synthesized a simulated spacer with 32nt at both ends and a BsaI site in the middle, and carried it on PUC57-Kan plasmid. After that, sgRNA suitable for endogenous CRISPR system was carried to the spacer in the middle of the repeat region, and the spacer was under the control of a lactose-induced promoter. The homologous arm for repair was carried to SmaI and Hind? of PUC57-Kan plasmid.

### Transformation and mutant screening

Plasmids used in this study were transferred to *Paracoccus-KDSPL-02* by electroporation. Then place in a constant temperature incubator at 37°C for incubation. The obtained incubation product were re-streaked onto LB plates supplemented with kanamycin (Kan) or chloramphenicol (Cm) to select for transformants. Plates were incubated for 72~84 h until colonies appeared.

This study need to re-streake incubation product onto LB plates supplemented with kanamycin (Kan) and IPTG for endogenous CRISPR/Cas system. After the colonies were observed, the desirable mutants were screened by colony PCR and then confirmed by Sanger sequencing. To cure the plasmid in the newly obtained mutant, successive transferring was carried out in LB liquid medium without any antibiotics. After 6–8 times transfer, the culture was then spread onto LB plates for colony development. The plasmid-cured mutant was obtained through PCR.

### Penicillin G degradation by whole *KDSPL-02* cells under visible light irradiation

A suspension solution (0.2 mL) of whole KDSPL-02 cells with a degradation activity of 2.0– 7.0 U was added to a 250-mL conical flask containing a 100-mL solution of 0.8–1.6 g L^−1^ penicillin G. The treatment was carried out at 120 rpm at 32 °C under visible light irradiation. Initially, persistent organic compounds were converted into more easily photodegradable compounds, thereby promoting photodegradation. The photocatalytic degradation of penicillin G was carried out in an annular reactor under visible light or LED irradiation with a medium mercury lamp (TQ 718 Z1 700 W) purchased from Heraeus at the reactor centre.

### Analytical methods

The changes of substrate and intermediate products of antibiotic penicillin G degradation were detected by HPLC. Penicillin G, Penicillium potassium thiazolate, and penicilloic acid concentrations in the fermentation broth were determined using high-performance liquid chromatography system equipped with UltimateR XB-C 18 (250 mm×4.6 mm,5 μm) and refractive index detector. The mobile phase was potassium dihydrogen phosphate (0.500 mol/ L, pH adjusted to 3.5 with phosphoric acid solution): deionized water: methanol =1:4:5. At a flow rate of 1.000 ml/min at 35°C. The ultraviolet detection wavelength is 225nm. Cell density was monitored using a cell density meter at 600 nm (OD_600_).

## DATA AVAILABILITY STATEMENT

Not applicable.

## AUTHOR CONTRIBUTIONS

PW conceived the study, carried out the experiments, and drafted the manuscript. CS and SX participated in the gene editing study. PW, CS, JM, SX, FC and JL assisted in the photoassisted biodegradation experiments. All authors read and approved the final manuscript.

